# Multienzyme deep learning models improve peptide *de novo* sequencing by mass spectrometry proteomics

**DOI:** 10.1101/2022.08.03.502594

**Authors:** Carlos Gueto-Tettay, Di Tang, Lotta Happonen, Moritz Heusel, Hamed Khakzad, Johan Malmström, Lars Malmström

## Abstract

Generating and analyzing overlapping peptides through multienzymatic digestion is an efficient procedure for *de novo* protein using from bottom-up mass spectrometry (MS). Despite improved instrumentation and software, *de novo* MS data analysis remains challenging. In recent years, deep learning models have represented a performance breakthrough. Incorporating that technology into *de novo* protein sequencing workflows require machine-learning models capable of handling highly diverse MS data. In this study, we analyzed the requirements for assembling such generalizable deep learning models by systematically varying the composition and size of the training set. We assessed the generated models’ performances using two test sets composed of peptides originating from the multienzyme digestion of samples from various species. The peptide recall values on the test sets showed that the deep learning models generated from a collection of highly N- and C-termini diverse peptides generalized 76% more over the termini-restricted ones. Moreover, expanding the training set’s size by adding peptides from the multienzymatic digestion with five proteases of several species samples led to a 2-3 fold generalizability gain. Furthermore, we tested the applicability of these multienzyme deep learning (MEM) models by fully *de novo* sequencing the heavy and light monomeric chains of five commercial antibodies (mAbs). MEM models extracted over 10000 matching and overlapped peptides across six different proteases mAb samples, achieving a 100% sequence coverage for 8 of the ten polypeptide chains. We foretell that the MEM models’ proven improvements to *de novo* analysis will positively impact several applications, such as analyzing samples of high complexity, unknown nature, or the peptidomics field.

## Introduction

Bottom-up mass spectrometry-based proteomics (MS) is focused on the sensitive identification and quantification of peptides and, thereby, proteins in arbitrarily complex samples[1,2]. In the standard workflow, peptides are first produced through the proteolysis of proteins with the enzyme trypsin. In the following step, the generated peptides are separated by liquid chromatography and measured by mass spectrometry in tandem (LC-MS/MS). Finally, the peptide-spectrum matches (PSM), the assignment of the peptide sequences to individual MS spectra, are produced using comprehensive compendia of reference protein sequences database[3].

Some of MS’s remarkable applications are in the infection medicine proteomics field, where it is employed to characterize the molecular mechanism behind invasive bacterial diseases[4–6], modeling host-pathogen interactions[7–13] and investigate systemic proteome changes[14–18]. The use of the trypsin protease is justified by its efficiency, stability, and specificity to cleave only at the C-terminal of the basic residues, arginine, and lysine[19]. However, its applicability is limited by the amino acid composition of the target proteins and the pH of the digestion solution[20,21]. Proteases other than trypsin, such as Elastase, Glu-C, Asp-N, Pepsin, ProAlanasa, are employed to achieve different cleavage patterns or work in various pH ranges[22–25]. Despite the increasing maturity of bottom-Up MS, peptide identification is restricted to the sequences included in a reference database. Consequently, it is unattainable to study proteins derived from organisms without sequence or which are extinct, environmental samples, and microbiomes. Other examples involve therapeutic monoclonal antibodies, i.e., immune system proteins composed of heavy (HC) and light (LC) chains containing conserved and variable regions. The latter region is typically not contained in the traditional sequence databases for either chain[24,26,27]. To overcome this limitation, *de novo* MS peptide sequencing is intended to extract partial or complete sequence information directly from collected MS spectra. In this strategy, the identities and positions of the amino acids are determined by the differences in mass of a series of consecutive fragments, for example, fragment ions of type *b* and *y*. To this end, programs have been created which implement algorithms based on graph theory, Hidden Markov models, linear and dynamic programming, such as PEAKS[28], NovoHMM[29], Lutefisk[30], Sherenga[31], pNOVO[32,33], and PepNovo[34], among others. As in other fields of proteomics[35], the application of deep learning represented a performance breakthrough in *de novo* MS peptide sequencing, as in the case of DeepNovo[27]. Deep learning algorithms attempt to simulate the behavior of the human brain—albeit by using many connected layers of neurons, which allows it to learn multiple levels of representation of high-dimensional data[35–38]. This key aspect translates into revolutionary advances in many research fields, such as image processing[39], speech recognition[40], and natural language processing[37]. In the supervised learning flavor, a model learns to make predictions based on labeled training data. Here, features like the amount of data and their diversity directly impact the resulting model’s generalizability, i.e., their ability to react to new data and make accurate predictions. Therefore, generalizability is central to the success of a model and its further implementation[36,38]. DeepNovo software outperformed other state-of-the-art methods at the level of amino acids and peptides. It combines convolutional and recurrent neural networks and local dynamic programming to learn the characteristics of tandem mass spectra, fragment ions, and sequence patterns of peptides. A later version (DeepNovoV2) added an order-invariant network architecture (T-Net) and a sinusoidal m/z positional embedding[41], which exceeds its predecessor by at least 13% at the peptide level[42].

It has been reported that the generation and analysis of overlapping peptides through multi enzymatic digestion is an efficient procedure for tandem MS *de novo* protein sequencing[24,25,33,43]. This approach can even resolve some of the challenges encountered in conventional strategies, which depend on the cloning/sequencing of coding mRNAs[43–45]. Given the mentioned facts, integrating DeepNovo deep learning architecture to handle the multi enzymatic MS samples can be game-changing for the *de novo* protein sequencing field. In order to accomplish this, it requires generalized models for successfully decoding the peptide sequences from MS samples treated with a wide variety of proteases. Previous DeepNovo studies reported models trained exclusively from a compendium of tryptic peptides. i.e., trypsin-SEM models[27,42]. This fact leaves the door open to questions related to the generalizability of the trypsin SEM models. Firstly, it is uncertain whether these models have extended applicability to other MS datasets, i.e., having high accuracy on samples generated using proteases with different cleavages specificities to the one employed to produce the model’s training set. In like matter, how the training set’s composition impacts the resulting model’s generalizability. Similarly, the effects of characteristics of the target spectra that facilitate peptide sequencing remain unexplored.

We studied the requirements for building generic DeepNovo models for the *de novo* MS sequencing task in the present work. For that purpose, we analyzed how the peptide composition and size of the training set affect the resulting model’s generalizability. The efficiency of these models was assessed by calculating the peptide recall on two highly sequence diverse test sets. Data showed reiteratively that using a collection of peptides with a wide variety of N- and C-termini amino acids led to 76% more generalizable models than the termini-restricted ones. Furthermore, DeepNovo models kept improving in the *de novo* peptide MS sequencing task as we continued extending the training set data with the multienzyme digestion of various species samples. We further proved the relevance of these multienzyme deep learning (MEM) models by *de novo* sequencing the heavy and light monomeric chains of five commercial monoclonal antibodies (mAbs). MEM models fully sequenced 8 of 10 target proteins, extracting over 10000 confirming and overlapping peptides from mAb MS samples digested with six different proteases. We consider that MEM models, combined with other mass spectrometric techniques, will help *de novo* analyze MS samples of higher complexity, such as the mixture of mAbs.

## Results and discussions

To integrate DeepNovo into the *de novo* protein sequencing pipeline, we need deep learning models capable of performing *de novo* sequencing in MS spectra of samples digested with numerous proteases. Therefore, it is first mandatory to determine the basis for building such generic models. For that purpose, we explored the effect of the training set composition on the resulting model generalizability, following the workflow in **Figure 1**. We initially created five peptide datasets by digesting Detroit 562 cell line samples with five proteases: trypsin, chymotrypsin, elastase, gluc, and pepsin (see **Material and Methods** section for LC-MS/MS and spectra annotation details). In each dataset, 21492 annotated spectra were randomly selected and split into training(90%), validation(5%), and test (5%) sets. We then systematically built multiple models from the training sets data. In order to assess all models’ generalizability, it was essential to evaluate their performance on a dataset composed of highly variable peptides in terms of amino acid composition and peptide length distribution. For that reason, we constructed the Detroit test set by merging all five test sets. Here, we used the peptide recall as a quantitative metric for the generalizability assessment. In addition, given that the protease employed during the sample preparation has a direct effect on the resulting peptides collection termini variability, we calculated the number of unique trimers on the N-terminal (Tn) and C-terminal (Tc) for all the generated models’ training sets in this study. Tn and Tc are quantitative metrics for the extent of the training sets’ variability at each peptide termini. Higher values of Tn and Tc represent higher variability in the peptide dataset at N and C-termini, respectively. We also introduced the diversity factor (DF), defined as log(Tn/Tc), as a measure of the variability balance between the training set’s N- and C-terminus. DF values near zero represent models with a better balance between the number of trimers at each terminal. Similarly, positive and negative DF values indicate a larger proportion of Tn and Tc, respectively.

**Figure 1.**
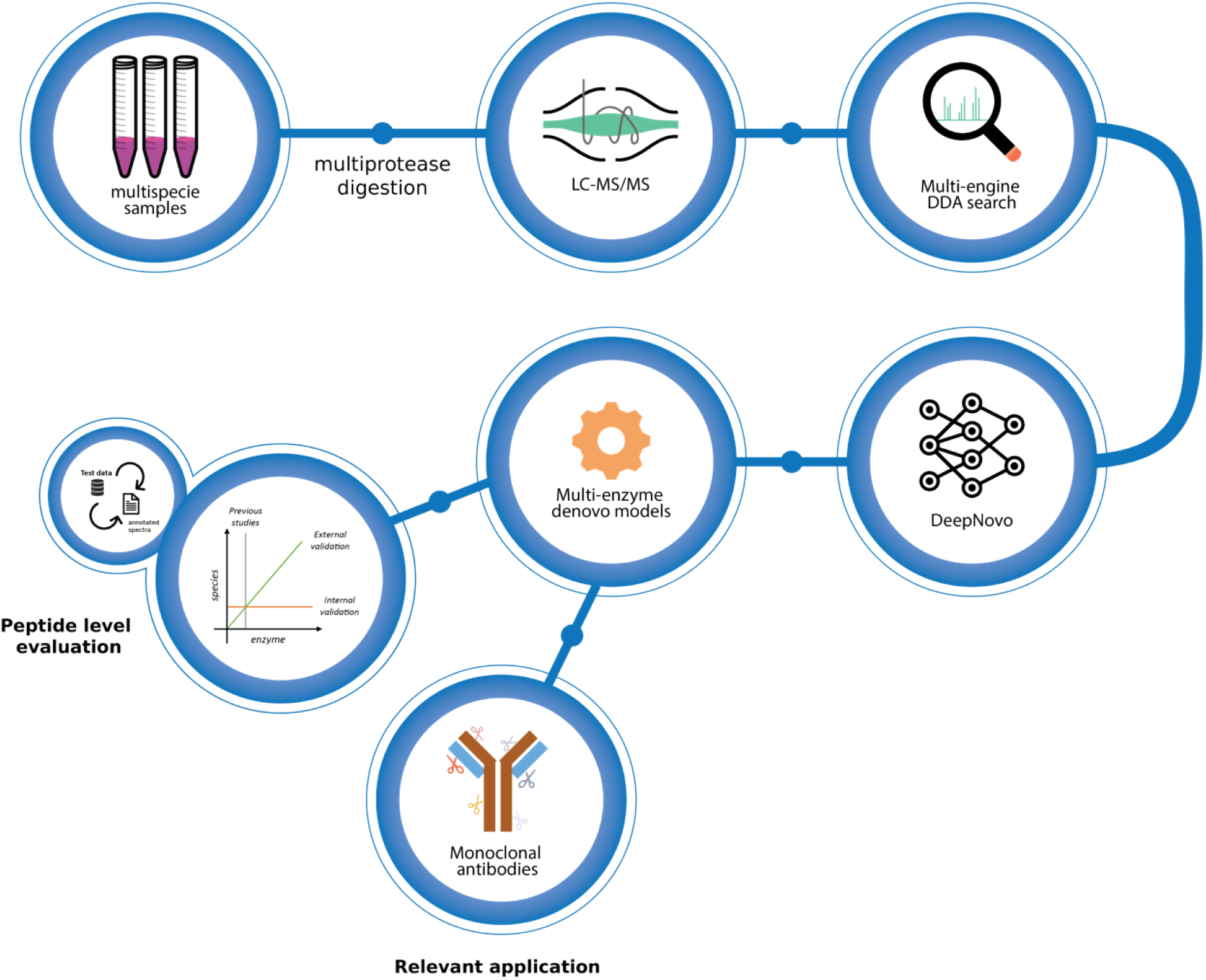
We started with three sample cohorts; Detroit 562 cells, 5 commercially available antibodies, and a large collection of samples from different species. The samples were aliquoted and digested using five enzymes, measured using LC-MS/MS, and analyzed using traditional database searches with multiple search engines. All data were also analyzed using the published DeepNovo deep learning model. Several DeepNovo models were created, see text for details, and evaluated in three ways. The internal validation evaluated the model performance on data generated with the same enzyme(s) as the model was trained with. The external validation evaluated the model performance using data generated with enzyme(s) different from the model creation data. We finally assessed each model’s performance in *de novo* sequencing five full-length antibodies.

### Nonspecific enzymes training datasets yield more generalized models

We built the first round of models from the five individual enzyme datasets and identified them as Single Enzyme Models (SEM). **Figure 2** displays the characteristics and performance of the SEM model on the Detroit test set. Two findings are worth mentioning regarding SEM models: 1) Using less specific proteases for the peptide generation leads to more N/C termini balanced training sets (**Figure 2A**). In contrast to pepsin, trypsin protease has a high specific cleave pattern that generates a training set with high Tn and low Tc values, as the peptides end in either arginine or lysine amino acids. This observation is supported by the DF values for SEM models, i.e., pepsin (0.52) < chymotrypsin (0.76) < elastase (0.79) < Glu-C (0.88) < trypsin (1.50); 2) Models’ generalizability correlates inversely with DF values (**Figure 2B**). Data shows that SEM models built with less specific enzymes, such as pepsin, chymotrypsin, and elastase, outperform 14 - 46% of those generated from using proteases with more specific cleave patterns, like gluc and trypsin, on nonspecific N/C termini peptides datasets. These differences in SEM models’ generalizability are explained when considering their performance on the Detroit set’s components (**Figure 2C**). We found that the most contributing factor was related to the models’ performance on inter-enzyme datasets, e.g., where the proteases for generating the training and test sets differed. For example, the pepsin-SEM model performed 46 - 86% better than the trypsin-SEM model on chymotrypsin, elastase, and gluc peptide datasets. In addition, all SEM models performed best when there was a match between the protease employed to generate the SEM model’s training set and the Detroit set’s portion. In these cases, peptide recall ranged from 0.46 to 0.69. These results are comparable to previous DeepNovo works where only trypsin was used[27,42]. Here, less specific SEM models outperformed 6 - 48% of the highly cleave pattern-specific ones. These results suggest that SEM models generated from the digestion with trypsin and gluc are more biased at the spectra decoding stage, especially for purposing the C-terminus peptide amino acids.

**Figure 2.**
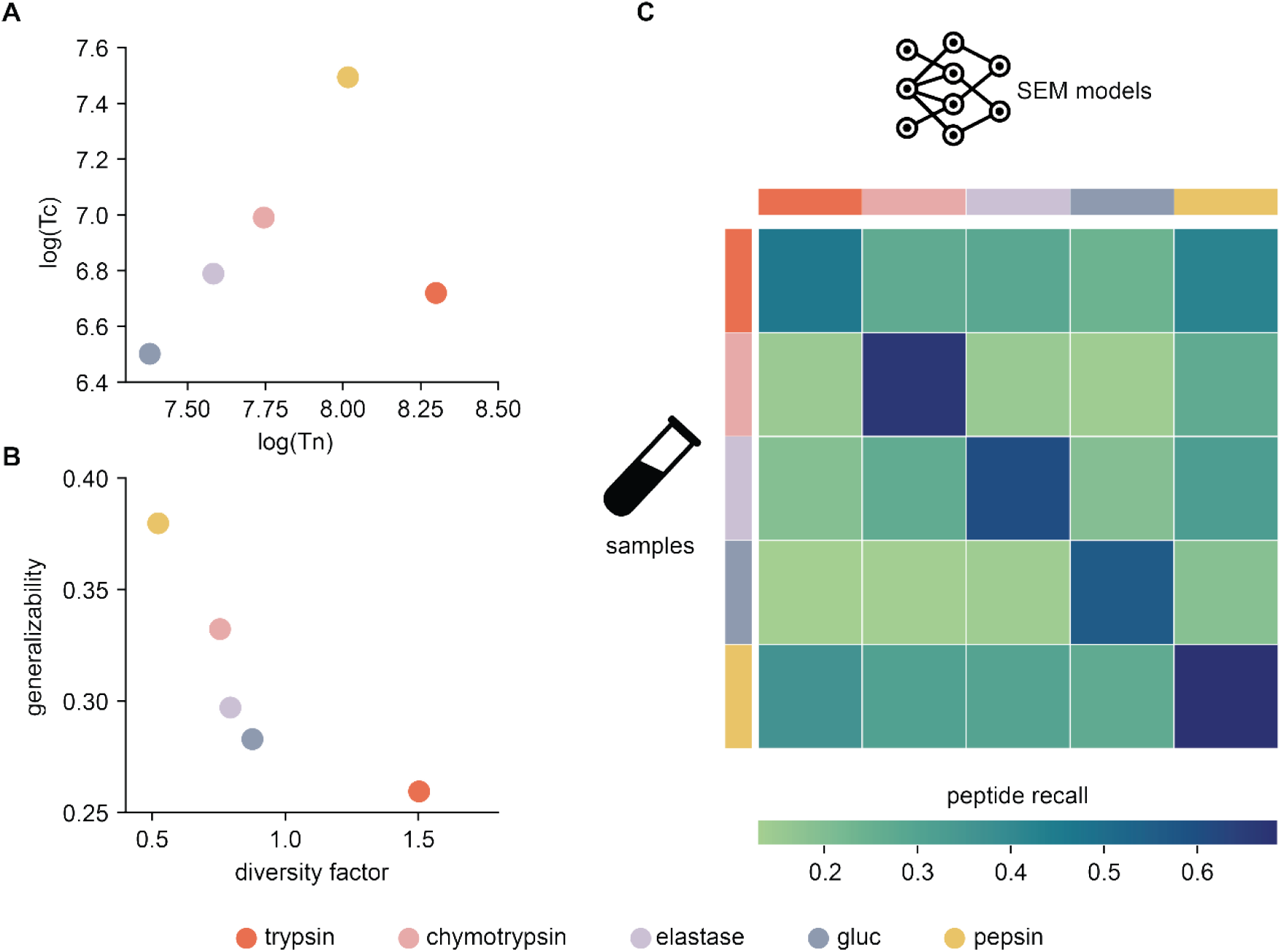
Single Enzyme models (SEM models) performance on Detroit test set: **A)** Tn and Tc values for SEM training sets; **B)** SEM models’ generalizability vs. training set’ diversity factor **C)** SEM models peptide performance on the individual enzyme-specific datasets composing the Detroit test set. The color scheme for both models and samples is at the bottom.

Inspired by the results of the first round, we then decided to test if it was possible to modulate models’ generalizability as a function of their training set’s diversity factor. Here, the number of spectra selected for the construction of each model was equal to those of SEM models for a fair comparison. For that purpose, we built new models distributed in two categories: 11 monoterminal (MoTM) and 12 multiterminal (MuTM) models. In MoTM models, all training set’s peptides share a specific amino acid at one termini position; for example, in the ThrN and PheC MoTM models, all peptides have a Thr or Phe amino acid at the N- or C-terminal, respectively. Contrary, MuTM models prioritized maximum variability at both terminals by selecting peptides from all SEM models’ training sets. **Figure 3** displays MoTM and MuTM model characteristics and performance on the Detroit test set.

**Figure 3.**
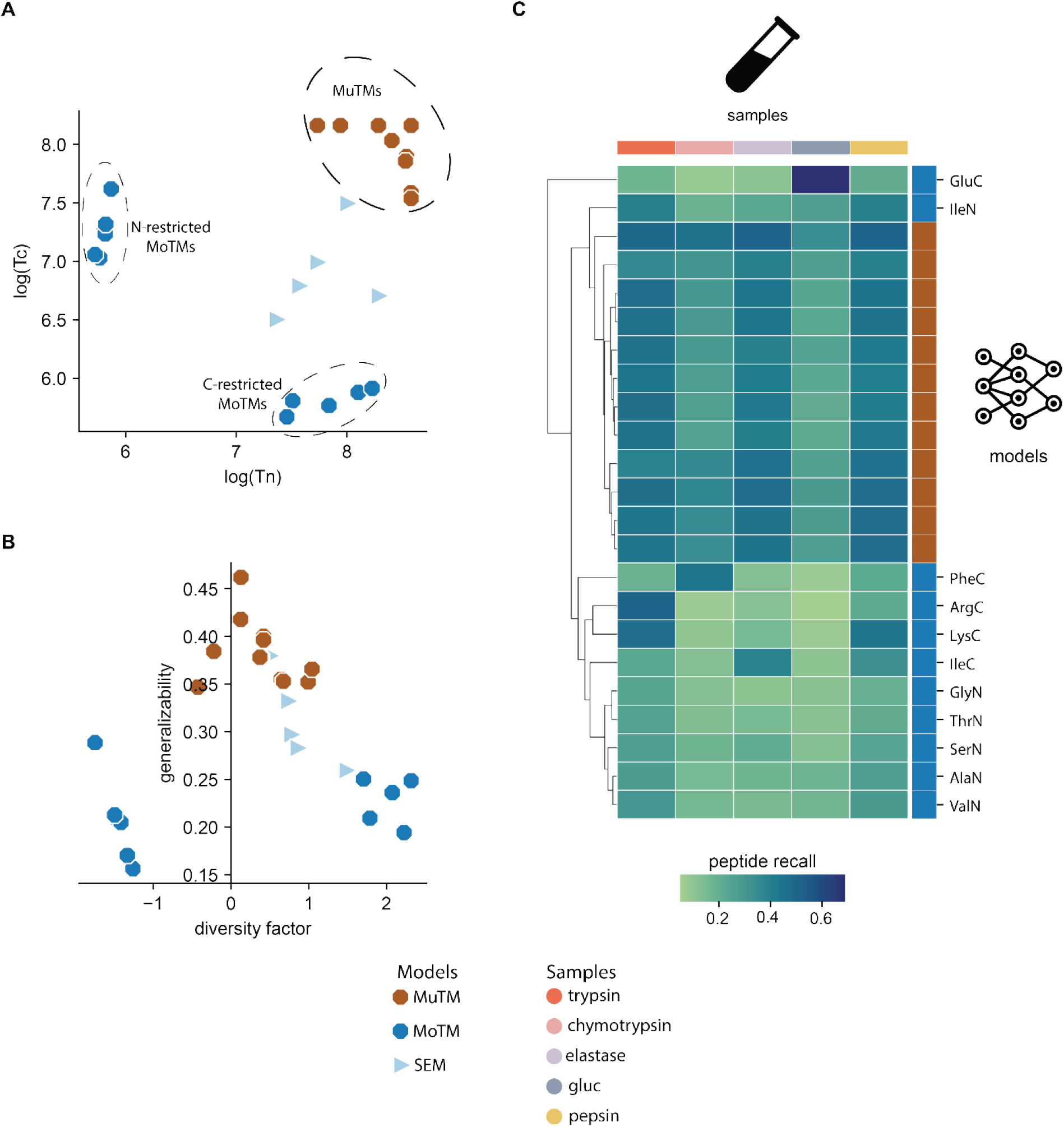
Modulating models’ generalizability by varying their training sets’ termini variability. Comparing Monoterminal (MoTM) and Multiterminal (MutM) models’ characteristics and performances on the Detroit test set: **A**. Training sets N/C-termini variability for MoTM and MuTM models; **B**. Models’ generalizability as a function of their termini diversity factor; **C**. MoTM and MuTM models peptide recall on Detroit test set components. We included SEM models’ Tn, Tc, and diversity factor parameters as a quantitative reference. Color and shape schemes for models and sample types are at the bottom.

Considering SEM models as reference, three new groups are distinguishable regarding Tn and Tc values distributions (**Figure 3A**). Two groups belong to MoTM models, which have low Tn and Tc values for the N-termini and C-termini restricted MoTM models, respectively. The third group belongs to the MuTM models, containing high values for both Tn and Tc parameters. **Figure 3B** shows that MuTM models are more termini-balanced and generalized than all MoTM ones. According to the mean values of the peptide recall on the Detroit test set, MuTM models outperform 76% of MoTM models. Moreover, half of the MuTM models generalize better than the pepsin-SEM model, while the other half of the models were better than the chymotrypsin-SEM one. In contrast, 10 of 11 MoTM-biased models were worst than the trypsin-SEM model at generalizing. On the other hand, the models’ performance on the Detroit test set’s components shows how MuTM models cluster together as they exhibited more uniform peptide recall values across all sample types (**Figure 3C**). On the contrary, the MoTM models’ performance depended on the cleave rules’ overlap between the sample and the model’s training set. For example, the ArgC-MoTM model performed best on the trypsin sample. However, peptide recall values dropped 57 - 90% in the remaining sample types. Other similar cases involved the GluC and PheC-MoTM models. These observations suggest that, under the same amount of training data, it is possible to design more generalizable models by maximizing and balancing the training set’s Tn and Tc values.

### Large multienzyme models perform best

Since all SEM models perform the best on similar data types as the model’s training set, we then decided to build 26 new models by mixing all possible combinations of the five Single Enzyme models’ training sets, e.g., multienzyme models (MEM) from the combination of 2 (n=10), 3 (n=10), 4 (n=5), and 5 (n=1) SEM-datasets. Here, the MEM model composed for all five Detroit 562 peptide datasets was called the Kilo MEM. Data shows that appending one or more different peptide datasets to any existing SEM dataset yields growth in Tn, Tc, and generalizability parameters for the resulting MEM model (**Figure 4**). As expected, the increase in Tn and Tc values was more noticeable when the merged datasets did not share the same cleave rules as in chymotrypsin - gluc and trypsin - elastase - gluc dataset combinations (**Figure 4A**). Furthermore, generalizability and diversity factor values suggest that MEM models generalize better and are more termini-balanced as we increase the number of peptide datasets (**Figure 4B**). An illustrative example of MEM models’ rising performance is shown in **Figure 4C**, where we displayed the path to generating the Kilo MEM from the pepsin-SEM model. Two observations are worth mentioning: 1) new datasets contributed positively to the resulting MEM model’s generalization, and 2) the formed MEM model always performed better than its antecessors models. The Kilo MEM model not only doubles the termini peptide dataset variability but also produced an increase of 38% in diversity factors concerning all SEM models. As a result, the Kilo MEM model outperforms 1.8 - 2.4 times the SEM models.

**Figure 4.**
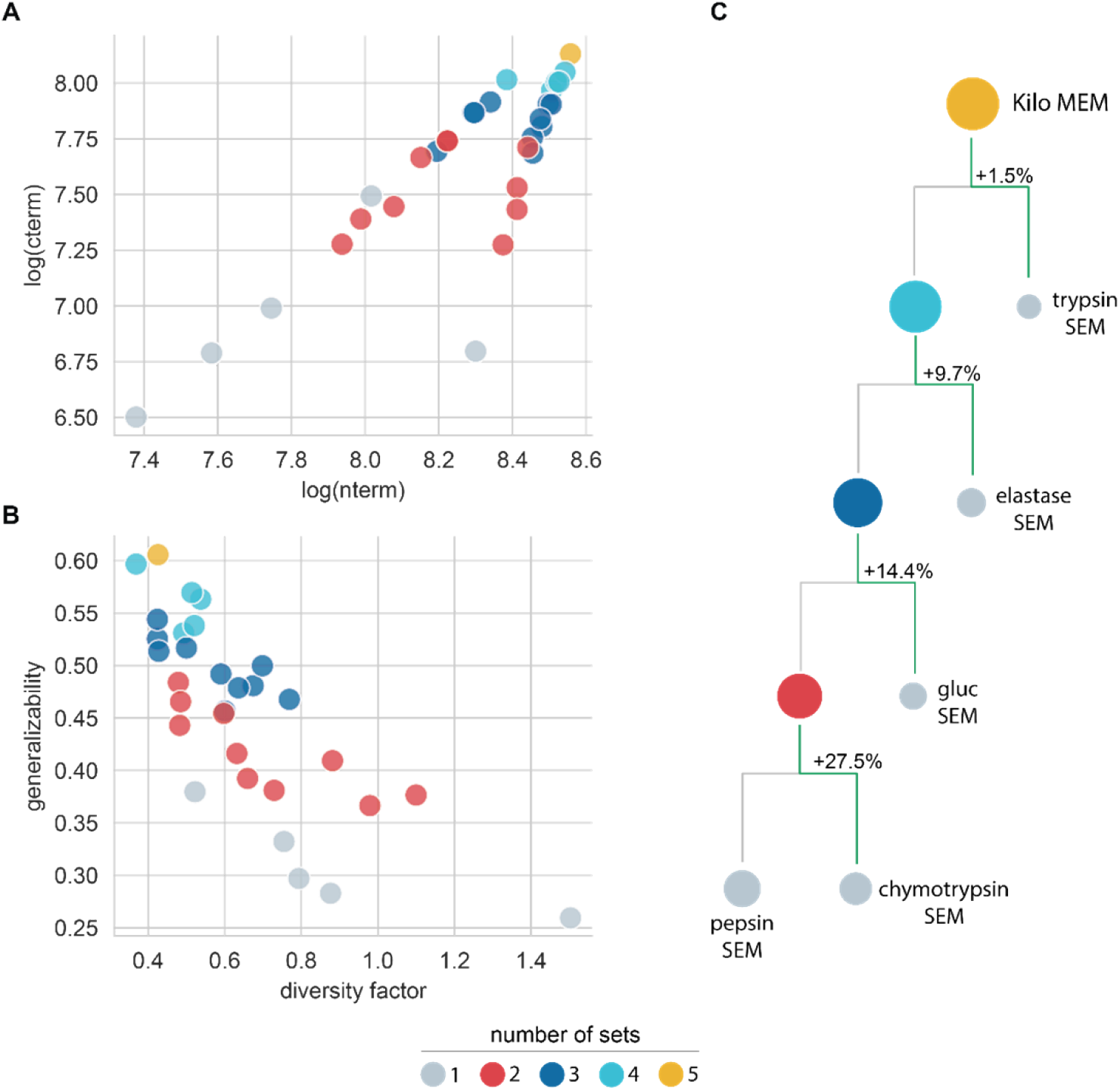
Characteristics and performance of multienzyme (MEM) models: **A**. Training sets’ Tn and Tc values; **B**. MEM models’ generalizability vs. diversity factor; **C**. Sequential building of Kilo MEM model from all five SEM datasets. The size of circles is proportional to their generalizability values on the Detroit test set. The color scheme at the bottom reflects the models’ characteristics variation with the number of combined datasets. SEM model data (displayed in gray) were used as reference.

The results of the SEM and MEM models demonstrated that features such as the training set’s size and peptide sequence variability significantly impact the resulting model’s generalizability. At this point, we hypothesized that expanding sequence variability by creating a training set that includes peptides across different species will lead to a more generic model than the Kilo MEM model. To prove it, we generated an external Giga dataset by digesting various species samples, such as *Saccharomyces cerevisiae, Escherichia Coli, Equus caballus, Streptococcus pyogenes*, and *Mus musculus* with trypsin, chymotrypsin, elastase, and gluc proteases. We followed the same protocol for sample injection, MS detection, and database search (See **Material and methods**). After spectra annotation, the Giga dataset was ten times larger than the Detroit 562 dataset. We then trained and applied the Giga MEM model to the Detroit test set and compared the results with the Kilo MEM model. Data shows that the Giga MEM model generalized 29.4% better than the Kilo MEM model, outperforming 24 - 41% in all Detroit test set’s sample types (**Figure 5**). In the same way, the Giga MEM model generalizes 2.1 - 3.0 times better than the SEM models.

**Figure 5.**
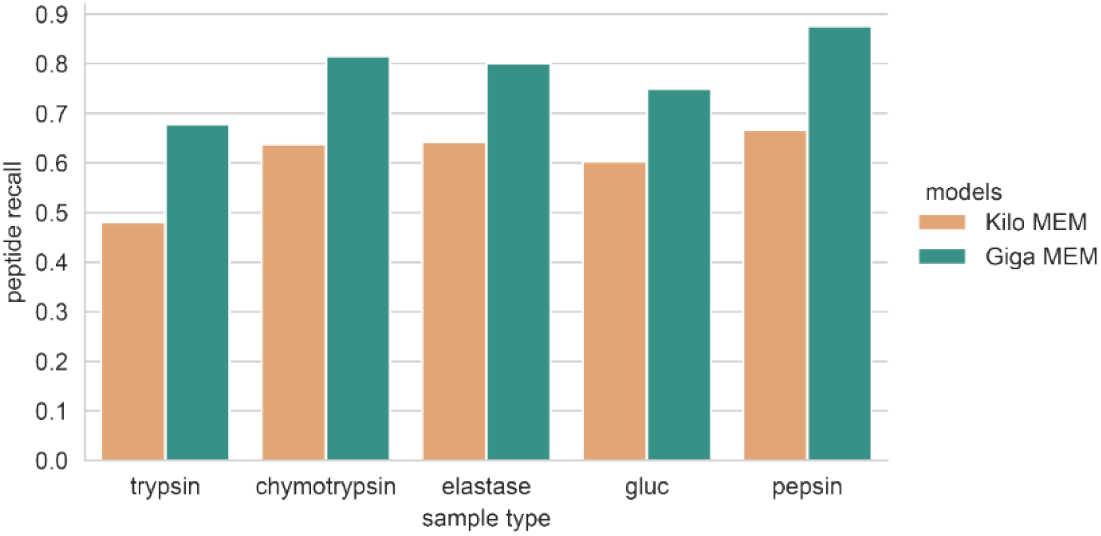
Comparing the performance of the Giga and Kilo MEM models across the Detroit test set’s different sample type components.

The Giga dataset was also used as an external test set. Specifically, we tested the generalizability of the 5 SEM and 26 MEM models. Interestingly, generalizability values on the Giga test set supported our previous findings on the best conditions to build more generic models (**Supplementary Information A**). Here, it is crucial to mention the pepsin-SEM model results; In the Detroit test set case, the most considerable portion of de novo sequenced spectra corresponded to peptides generated with the same protease as the SEM model’s training set. However, pepsin was not part of the multienzyme protocol for generating the Giga external peptide test set. Despite that, the pepsin-SEM performed best among all SEM models. Overall, generalizability results on the Detroit and Giga test sets suggest that, like other deep learning architectures, DeepNovo kept improving in the *de novo* peptide MS sequencing task as we fed the model with extensive and highly diverse peptide MS data.

### Fragment ions distribution impact MS *de novo* peptide sequencing

After establishing the criteria for building generalizable models, we further explored how the peptide composition impacts the ability to *de novo* sequence its spectrum correctly. In this respect, we studied the Kilo MEM model results on the Giga test set (**Figure 6**). Initially, we evaluated the effect of the peptide length distribution on the overall deep learning model’s performance by tracking the peptide recall as we varied the maximum peptide length (**Figure 6A**). We observed that performance decreased as we included longer peptides in the test set. Data shows that the probability of *de novo* MS sequencing correctly 6-residue peptides was 86.1% and fell quickly to 40% when considering peptides of up to 14 residues.

**Figure 6.**
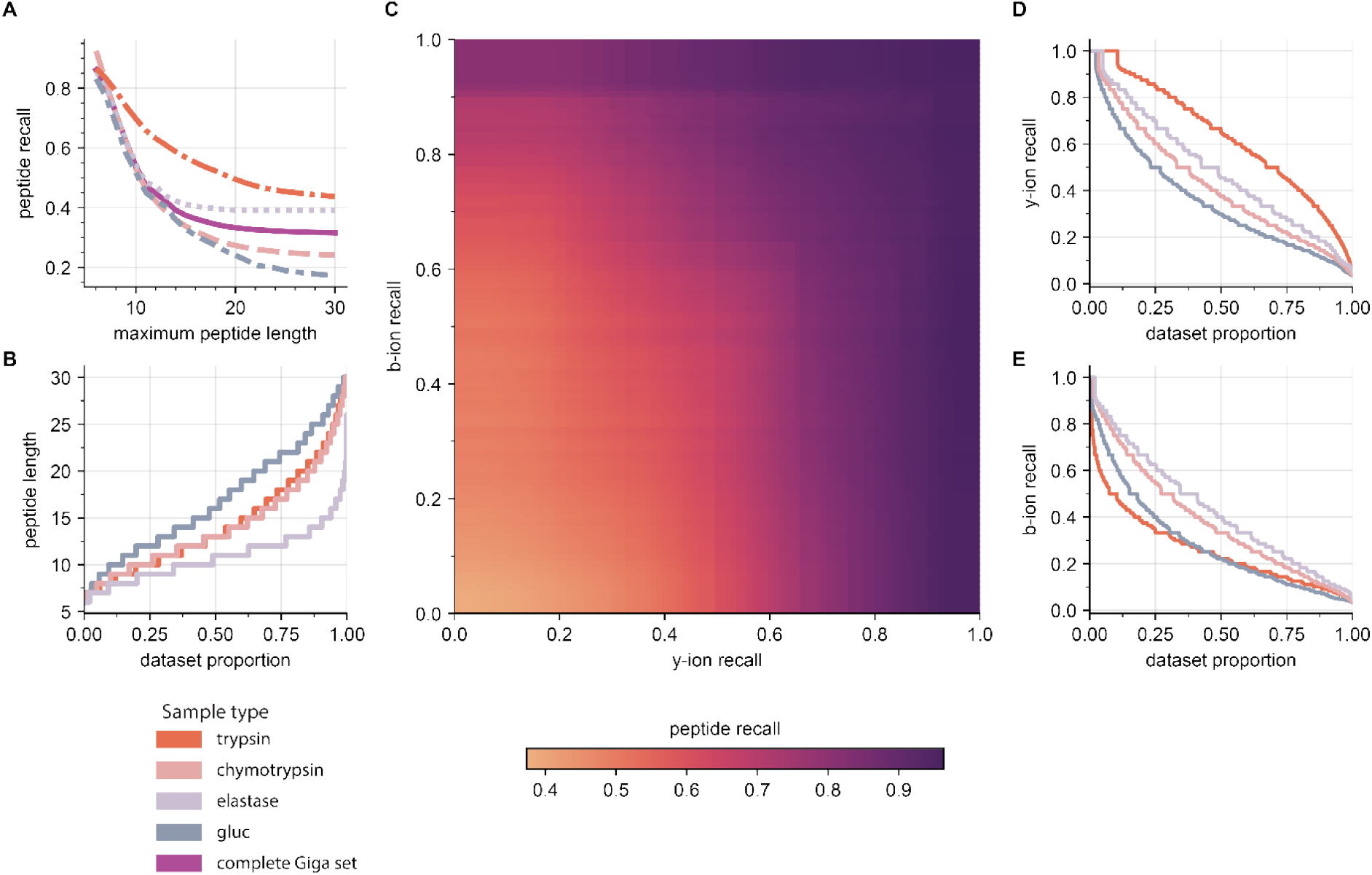
Kilo MEM model *de novo* sequencing results on the Giga external test set: **A**. peptide recall as a function of the maximum peptide length in the test set; **B**. peptide length distribution for all the Giga test set’s sample types; **C**. peptide recall as a function of the minimum singly-charged b-y ion recall grid values; D. y-ion recall and **E**. b-ion recall distributions for the trypsin, chymotrypsin, elastase, and gluc sample types spectra. The color scheme for the sample types is at the bottom.

Moreover, this performance decay differed for all components of the Giga set, suggesting that the identity of the peptides also impacts their chance of being MS sequenced. To explain these differences across the four datasets, we calculated the peptide length distribution (**Figure 6B**). Data shows that 75% of data in the elastase dataset are peptides of length 12 or shorter, explaining why it was more accessible to *de novo* MS sequence elastase data over chymotrypsin and gluc data. For the latter, 75% of the data were peptides of length 13 or longer.

Since the peptide length distributions could not explain performance differences related to the trypsin sample, we further calculated singly-charged b- and y-ion recall for all peptides spectra composing the Giga test set, e.g., the proportion of the fragment ions found experimentally over the total expected ones theoretically. Here the ion recall is a quantitative metric of the ability of a particular peptide to produce b/y-ions under specific experimental conditions[24,25,46,47]. For the fragment ions extraction, the m/z tolerance was 15ppm. We also calculated the peptide recall as a function of the minimum values for the b/y-ion recall pairs.

The b/y-ions recall grid shows that the probability of *de novo* MS sequence correctly a peptide increase with its capacity of producing either b- or y-ions (**Figure 6C**). Data shows that the peptide recall was higher than 70% when peptides produced at least 80% and 60% of the expected b- and/or y-ion fragments. These results suggest that the *de novo* MS sequencing performance on a specific sample type is bound to its b/y ion recall distributions. **Figure 6D** shows that the y-ion recall distribution order fits the peptide recall behavior for all sample types. It is worth mentioning that the tryptic peptides had the highest proportion of the expected singly-charged y-ions compared to the other sample types, explaining its remarkable performance across a wide range of peptide lengths (**Figure 6A**), e.g., 55% of the annotated spectra had at least 60% of the *y-*ions expected. For these peptides, y-ion fragments bear a charged residue, like arginine or lysine, which are more abundant and produce more intense peaks under the HCD fragmentation method [48,49]. On the contrary, the peptides from the digestion with gluc had a low proportion of y- and b-ions (**Figure 6E**). Furthermore, elastase b/y-ion recall distributions are consistent with a high proportion of short peptides.

### MEM models for full-length *de novo* sequencing of antibodies

Once we established the requirements for building generalizable models and how the quality of the input spectra impacts the subsequent *de novo* MS peptide sequencing process, we tested the efficiency of using the MEM models in the *de novo* protein sequencing pipeline. For this effort, we selected a challenging and biological interest system, such as the complete sequencing of monoclonal antibodies (mAbs). We aimed to fully *de novo* MS sequence the heavy (HC) and light (LH) chains of five commercial mAbs: Erbitux, Herceptin, Prolia, Silulite, and Xolair. We digested each mAb sample with six proteases: trypsin, chymotrypsin, elastase, gluc, pepsin, and aspn. It is worth mentioning that the latter enzyme was not part of the models’ generation protocol. On the other hand, we created the Giga+ MEM model by combining the training sets of the Kilo and Giga MEM models. We considered eight models (5 SEM + 3 MEM models) for comparison purposes. For analyzing results, we initially calculated the relative coverage for the entire variables space, i.e., models x samples x chains matrix (**Figure 7A**). This way, we got an insight into the model performance across all sample types and which enzymes facilitate the *de novo* sequencing of the HC and LC subunits. In addition, we examined the length distribution of the sequence matching peptides for all sample types (**Figure 7B**). These plots provide information about the decoding power of the models. It also shows the capacity of the different proteases to produce easily detectable peptides from the *de novo* MS sequencing perspective.

**Figure 7.**
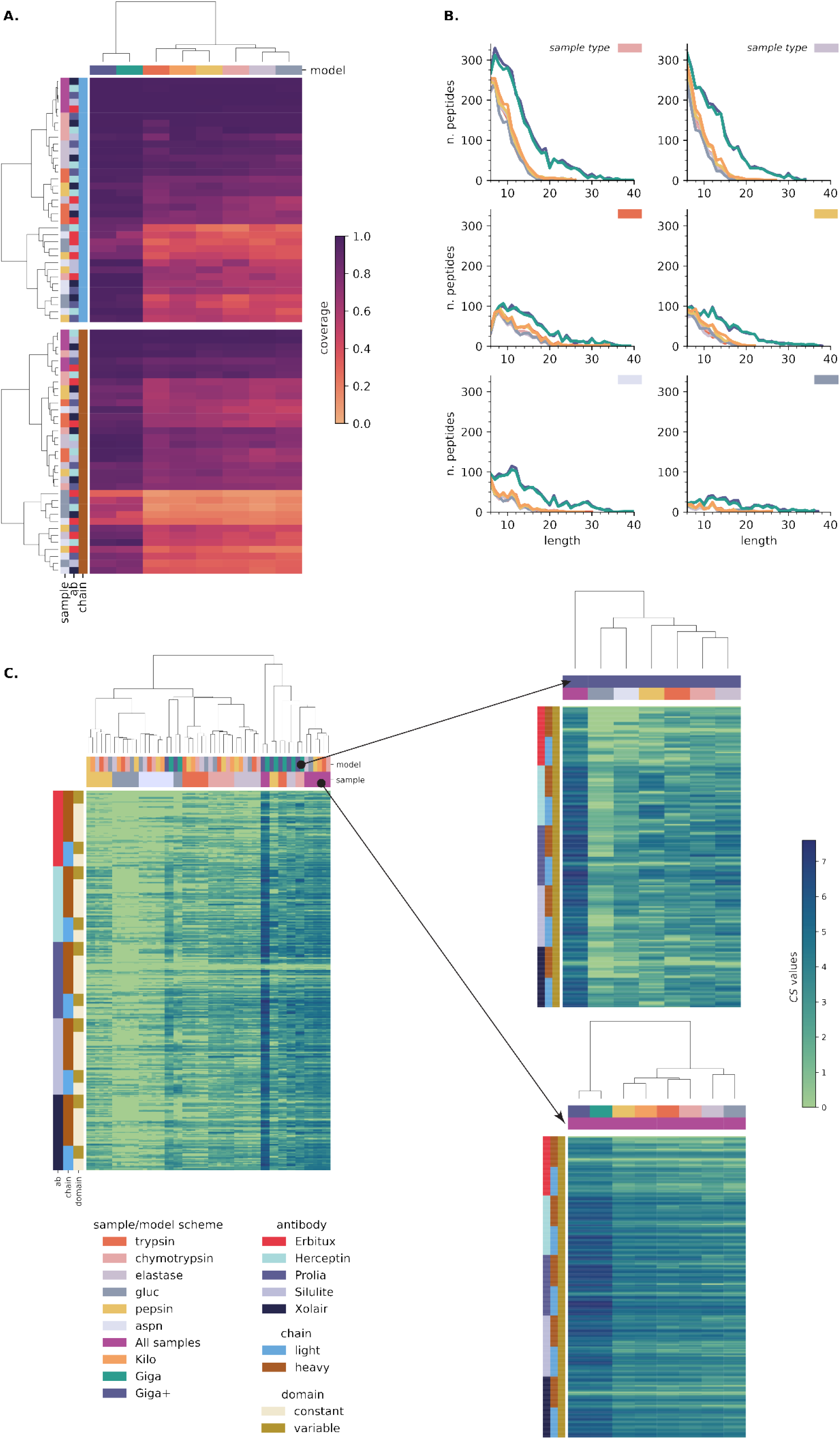
Complete de novo sequencing of commercial monoclonal antibodies by deep learning models: A) coverage of all the light (top) and heavy (bottom) chains for all types of samples. B) length distribution for all matching peptides extracted from each type of sample C) ***CS*** values for the variable and constant domains of monoclonal antibodies.

Regarding the sample types, the data shows that working with the chymotrypsin and elastase proteases had many benefits related to good protein coverage (**Figure 7A**) and the extraction of a high amount of matching peptides (**Figure 7B**). Data shows that digesting the samples with these proteases yields better individual protein coverage, wherein in 75% of cases, the sequence relative coverage was at least 0.80 and 0.75 for chymotrypsin and elastase, respectively. Additionally, the total amount of peptides extracted was 2 to 8 times greater than the rest of the proteases (**supporting information B**). It is worth noting that these were the only enzymes where, for lengths between 6 and 9, all of the considered deep learning models identified more than 100 peptides. These observations suggest that working with the chymotrypsin and elastase proteases leads to high amounts of readily *de novo* MS sequenceable peptides. As expected, the gluc and aspn digested samples got the lowest matching peptide extraction values, yielding the worst individual protein coverages. These proteases produced long peptides with low b- and y-ion recalls, making them more difficult to *de novo* sequence.

When comparing the performance of the deep learning models, the Giga and Giga+ MEM models were evident superior after considering the values of the protein coverages and the amount of matching extracted peptides parameters. For the Giga+ MEM model, the median value of protein coverage was 0.96 after considering all mAbs and sample types. Moreover, it extracted 10367 unique and confirming peptides, an amount 2 – 2.8 times greater than the Kilo MEM and all SEM models (**supporting information B**). Interestingly, and based on the same parameters, the pepsin SEM model was among the five SEM models. These findings supported our previous statements about the necessary criteria for building generalizable models. It is worth noting that the Giga+ MEM model sequenced all light chains and 3 of 5 mAbs heavy subunits for the combined sample results, i.e., Herceptin, Silulite, and Xolair mAbs. The remaining proteins had coverage of at least 0.97. It is essential to consider that, in mAb, the HC subunit can bear glycans in their constant region [50,51]. In some cases, such as for Erbitux, glycans are also found in the HC variable region [52].

As the overlapping of peptides is necessary for the assembly of protein sequences, we also decided to go deeper into the analysis of MAbs de novo results and introduce the *confident positional score (CS)*. For a residue in the position *i* of the protein sequence, is defined as *C*_*i*_ = *log*_2_(*f*_*i*_ + 1). Here *f*_*i*_ is the positional frequency for position *i*, i.e., the number of *de novo* sequenced matching peptides for position *i* in the protein sequence (**Figure 7C**). Higher consecutive *CS* values represent regions with more evidence in the *de novo* protein sequencing process, being especially important for MAbs HC and LC variable regions, for which the sequences are unknown. In contrast, sequence regions with no detected peptides have a zero positional frequency, ergo, a zero CS value. After combing all sample types, the Giga+ MEM model got a positional frequency greater than ten for 90.7% of the amino acids comprising the study mAbs. Moreover, this parameter value increased to 50 or more for 45.7% of said amino acids. Similarly, there were no confirming peptides for only 0.03% of residues. Furthermore, For the mAbs variable region, the median positional frequency was 45 and 51 for the HC and LC subunits, respectively (**Supplementary Information C**). For the five HC subunits, data show that CS values decreased up to 30% in the glycans’ surrounding regions, likely because of a steric effect as these bulky species prevent efficient digestion. In the case of the Erbitux mAb, the regions with zero CS values matched the glycans location for the HC constant and variable domains (**Figure 8**), suggesting that removing the glycans should be incorporated in the sample preparation to guarantee the complete MS sequencing of mAbs. Given the coverage and positional frequency results, the findings discussed here set a precedent for using multienzymatic deep learning models as an alternative for sequencing proteins from their multienzymatic digestion.

**Figure 8.**
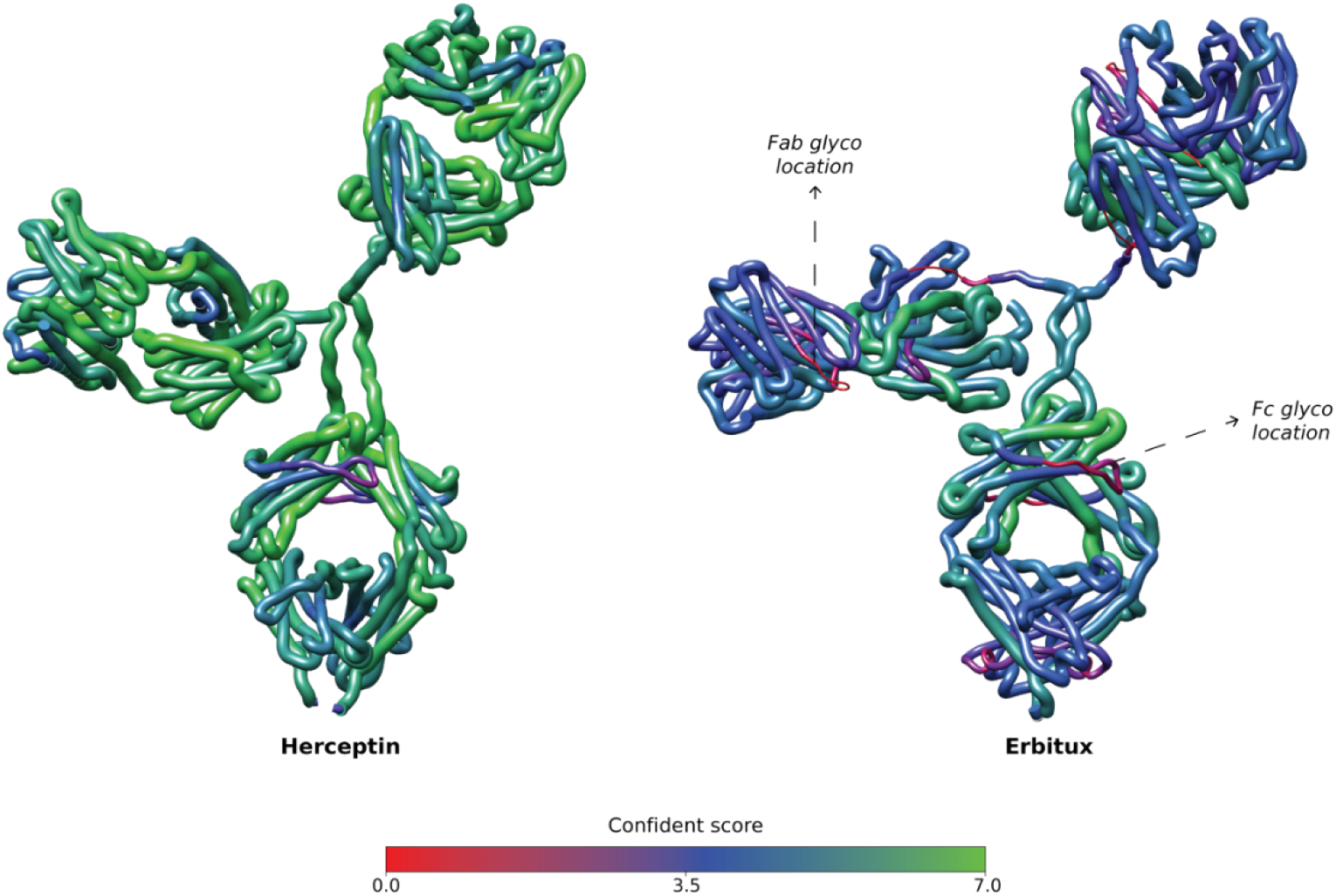
3D segmented worm representation of the mAbs with the highest (Herceptin) and lowest (Erbitux) *de novo* MS sequencing CS values. The thickness and color of the protein chains are proportional to their CS values.

In future studies, it may be interesting to explore using the multienzyme de *novo* sequencing protocol in conjunction with other complementary MS techniques like Top-down to sequence mixtures of mAbs.

## Conclusions

We proposed the use of MEM models to improve the *de novo* sequencing of peptides and proteins from DDA-MS data. Toward that aim, the effects of the properties of the training and test sets on the *de novo* sequencing process were explored. On the one hand, the data suggest that variability at both terminals, among the peptides which make up the training set, affects eventual generalizability. Consequently, the use of multiple proteases is recommended to generate more robust models. In the same vein, since DeepNovoV2 learns characteristics of spectra and sequences, an increase in the number of data points also improves the resultant model performance. These claims are supported by the peptide recall results for the test sets and the number of peptides extracted from the samples produced by multienzymatic digestion of commercial antibodies. On the other hand, it was discovered that the models’ *de novo* sequencing capacity is limited by the identity of the peptides and experimental conditions, which have direct consequences on their ability to produce ionic fragments of interest. This result explains why peptide recall fell with an increase in length of peptides, as well as the differences found among the samples from trypsin, chymotrypsin, elastase, and gluc. Finally, the findings described here will assist in other areas of peptidomics, the creation of Data-Independent-Acquisition libraries, and the sequencing of complex mixtures of monoclonal antibodies.

## Material and methods

### Sample preparation for mass spectrometry

#### Commercial antibodies

For sample preparation for mass spectrometry of commercial antibodies, 10 μg of each (Xolair, Novartis; Herceptin, Roche; SiLuLite, Sigma MSQC4 Universal Antibody Standard; Prolia, Amgen; and Erbitux, Merck) was denatured with 8M urea - 100 mM ammonium bicarbonate, the disulphide bonds reduced with 5 mM Tris (2-carboxyethyl) phosphine hydrochloride (TCEP) for 60 min at 37°C, 800 rpm, and alkylated with 10 mM iodoacetamide for 30 min in the dark at room temperature. The samples were diluted to a urea concentration <1.5 M with 100 mM ammonium bicarbonate. The antibodies were digested separately with 1 μg of trypsin, Promega; chymotrypsin, Promega; LysC/trypsin, Promega; elastase, Promega; GluC, Promega; or AspN, Promega for 18h at 37°C, 800 rpm. The digested samples were acidified with 10% formic acid to a pH of 3.0. The peptides were purified and desalted using SOLAμ reverse phase extraction plates (Thermo Scientific) according to the manufacturer’s instructions. Peptides were dried in a speedvac and reconstituted in 2% acetonitrile, 0.2% formic acid prior to mass spectrometric analyses.

#### Detroit 562 cell line

Briefly, ca.5 million cultured mammalian epithelial cells (Detroit 562 cell line) were kindly provided by Sounak Chowdhury. Suspension cells were first centrifuged at 5000g rcf, 4°C for 10 mins, followed by aspiration of supernatant and one time cold 1X PBS wash. Remained cell pellets were then added with 1 ml lysis working solution, composed of 1X RIPA lysis and extraction buffer, ThermoFisher, and 1X protease/phosphatase inhibitor cocktail, ThermoFisher. After 15 min incubation on ice, the cell lysates were precipitated by trichloroacetic acid (TCA), washed with 3X acetone, and dried in a speedvac. Completely dried protein extracts were reconstituted in 100 mM ammonium bicarbonate buffer and measured for protein concentration by BCA assays, ThermoFisher. 10 ug of cell lysate proteins were aliquoted for each reaction, 10 experimental replicates for each enzyme, 5 enzymes in total. Sample preparation of reduction, alkylation, enzyme digestion, acidification was described as above. Specifically, C18 spin columns were used for purification and desalting after the digestion. Except for the pepsin-digested group, LysC/Trypsin was introduced for a 1 hr pre-digestion prior to a 1 hr pepsin digestion.

### Liquid chromatography tandem mass spectrometry

The peptides of the digested commercial antibodies were analyzed on Q Exactive HF-X mass spectrometer (Thermo Scientific) connected to an EASY-nLC 1200 ultra-high-performance liquid chromatography system (Thermo Scientific). The peptides were loaded onto an Acclaim PepMap 100 (75μm x 2 cm) C18 (3 μm, 100 Å) pre-column and separated on an EASY-Spray column (Thermo Scientific; ID 75μm x 50 cm, column temperature 45°C) operated at a constant pressure of 800 bar. A linear gradient from 3 to 38% of 80% acetonitrile in aqueous 0.1% formic acid was run for 120 min at a flow rate of 350 nl min-1. One full MS scan (resolution 120 000 @ 200 m/z; mass range 350 – 1650 m/z) was followed by MS/MS scans (resolution 15000 @ 200 m/z) of the 15 most abundant ion signals. The isolation width window for the precursor ions was 1.3 m/z, they were fragmented using higher-energy collisional-induced dissociation (HCD) at a normalized collision energy of 28. Charge state screening was enabled, and precursor ions with unknown charge states and a charge state of 1, and over 6 were rejected. Data was additionally collected for non-tryptic digestions as above, but including peptides with a charge state of 1. The dynamic exclusion window was 10 s. The automatic gain control was set to 3e6 and 1e5 for MS and MS/MS with ion accumulation times of 45 ms and 30 ms, respectively.

### Computational analyses

#### Spectra annotation

A snakemake[53] was created for the DDA search. All DDA raw files were initially converted to Mascot generic format (MGF) by ThermoRawFileParser software (Hulstaert et al., 2020). Ursgal package[54,55] was used as an interface for searching the spectra against data’s Uniprot reference proteome using five search engines, namely MSGFPlus[56] (version 2019.07.03), MS Amanda[57] (version 2.0.0.17442), Comet[58–60] (version 2019.01.rev5), X! Tandem[60] (version alanine), and OMSSA[61] (version 2.1.9). Optional Met Oxidation (UniMod: 35), along with the fixed Cys carbamidomethylation (UniMod: 4) modifications, were considered in this study. Individual engine results were validated by percolator[62] (version 3.4.0), while the Combine FDR algorithm was implemented for combining results from all search engines[63]. Moreover, a threshold of 1% peptide FDR was set for decisive candidate inclusion.

#### De novo model generation and evaluation

The process of creating a model involves 3 steps, namely: 1) establishing the training, validation, and test sets; 2) creation of the input files for DeepNovoV2[42]; and 3) model training. For each one of the 5 DDA search results over the DetroitDetroit 562 data set digested with one specific protease, a total of 21492 annotated scans were selected. These were then randomly divided into training, validation, and test sets in proportions of 90%, 5%, and 5%, respectively. For the second step, a snakemake workflow was created for the extraction of the selected spectra and generation of the features and MGF files. Finally, model training was done in 20 epochs[27,42]. Maximum peptide length and mass were adjusted to 4000 Da and 30, respectively. These models were called SEM models.

The evaluation of the initial models was accomplished through full cross-validation. This was done with the aim of obtaining a perspective on the performance of each test set, as well as overall. The same modifications employed in database search were considered for all of the *de novo* searches in this research. Additionally, the maximum deviation of the precursor mass was adjusted to 15ppm. Peptide recall was used as a measure of the quality of the models.

#### Structural modeling

The Fc and Fab domains of each antibody were de-novo modeled separately by AlphaFold2[64,65], considering MMseqs2[66] to generate the multiple sequence alignment and homo-oligomer state of 1:1. For each selected model, the sidechains and the disulfide bridges were adjusted and relaxed using Rosetta relax protocol[67]. The loops in the hinge region were then re-modeled and characterized using DaReUS-Loop web server[68]. Finally, the full-length structure was relaxed, and all disulfide bridges (specifically in the hinge region) were adjusted using the Rosetta relax protocol. Visualization of the monoclonal antibodies was done through USCF Chimera software[69].

## Acknowledgment

This work was supported by Foundation of Knut and Alice Wallenberg (2016.0023).

